# Glisson’s capsule structure and function is altered in cirrhotic patients irrespective of etiology

**DOI:** 10.1101/2022.08.28.505570

**Authors:** Jessica Llewellyn, Caterina Fede, Abigail E. Loneker, Chet S. Friday, Michael W. Hast, Neil D. Theise, Emma E. Furth, Maria Guido, Carla Stecco, Rebecca G. Wells

## Abstract

**Background and Aims:** Glisson’s capsule is the interstitial connective tissue that surrounds the liver. As part of its normal physiology, it withstands significant daily changes in liver size. The pathophysiology of the capsule in disease is not well understood. The aim of this study was to characterize the changes in capsule matrix, cellular composition, and mechanical properties that occur in liver disease and to determine whether these correlate with disease severity or etiology.

**Methods:** 10 control, 6 steatotic, 7 moderately fibrotic and 37 cirrhotic patient samples were collected from autopsies, intraoperative biopsies and liver explants. Matrix proteins and cell markers were assessed by staining and second harmonic generation imaging. Mechanical tensile testing was performed on a test frame.

**Results:** Capsule thickness was significantly increased in cirrhotic samples compared to normal controls irrespective of disease etiology (69.62 ± 9.99 and 171.269 ± 16.65 µm respectively), whereas steatosis and moderate fibrosis had no effect on thickness (62.15 ± 4.97 µm). Changes in cirrhosis included an increase in cell number (fibroblasts, vascular cells, infiltrating immune cells and biliary epithelial cells). Key matrix components (collagens 1 and 3, hyaluronan, versican and elastin) were all deposited in the lower capsule although only the relative amounts per area of hyaluronan and versican were increased. Organizational features including crimping and alignment of collagen fibers were also altered in cirrhosis. Unexpectedly, capsules from cirrhotic livers had decreased resistance to loading in comparison to controls.

**Conclusions:** The liver capsule, like the parenchyma, is an active site of disease, demonstrating changes in matrix and cell composition as well as mechanical properties.

**Lay summary:** We assessed the changes in composition and response to stretching of the liver outer sheath, the capsule, in human liver disease. We find an increase in key structural components and numbers of cells as well as a change in matrix organization of the capsule in the later stages of disease. This allows the diseased capsule to stretch more under any given force, suggesting it is less stiff than healthy tissue.

**Graphical abstract:** 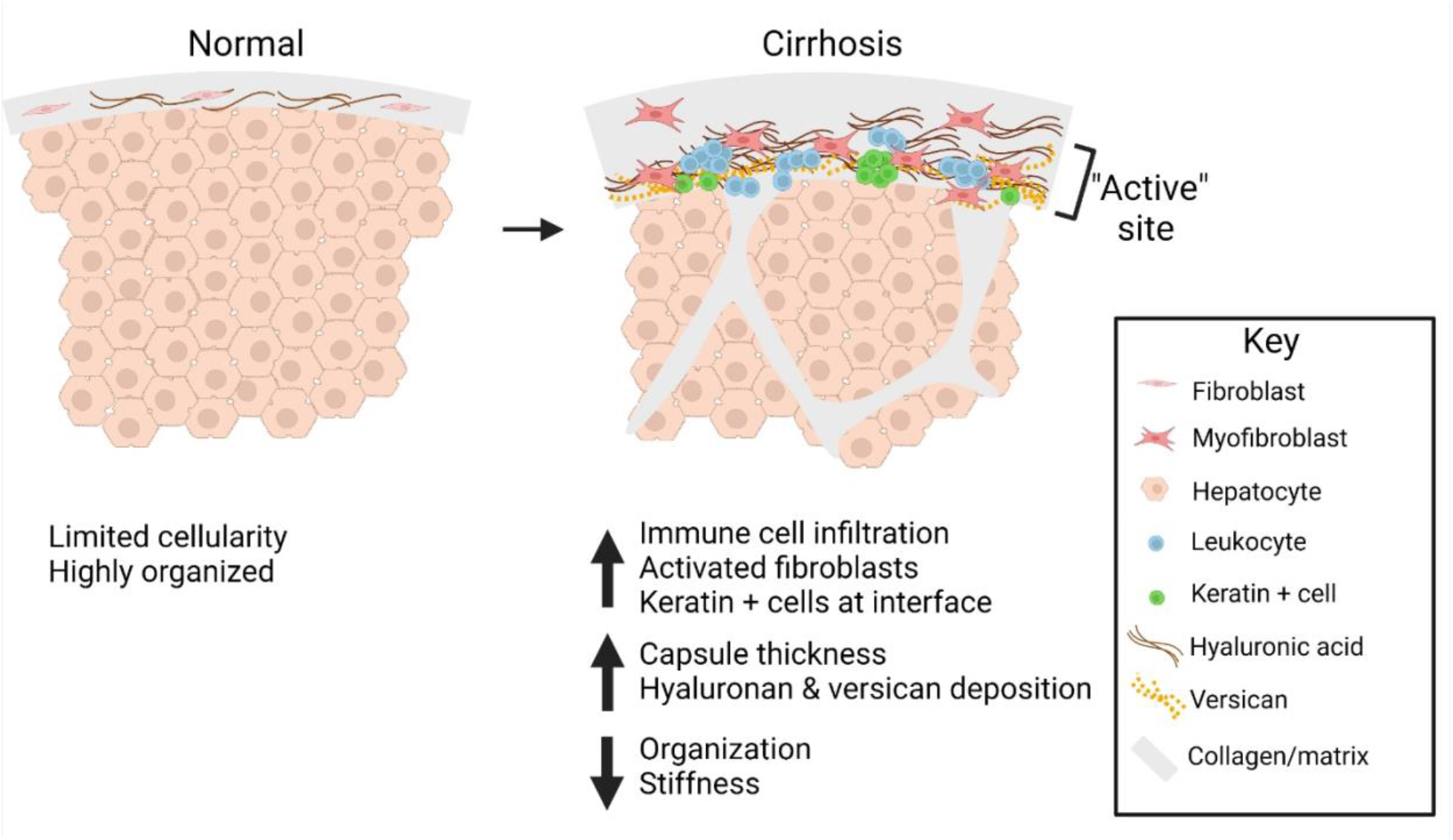

**Highlights:**

- The capsule is an active site of disease: thickness and cellularity increase markedly in cirrhosis
- Extracellular matrix composition and organization change in cirrhosis
- The cirrhotic capsule stretches more and is less stiff

## Introduction

The liver capsule, also called Glisson’s Capsule, is a layer of interstitial tissue that can also be classed as visceral fascia (1-3). It surrounds the liver and is continuous with the interstitial spaces and matrix surrounding the portal triads, likely playing a role in fluid exchange as well as cell migration within the hepato-biliary system (4, 5). The capsule has an important structural role, enclosing the parenchyma and protecting the liver from physical impact. Additionally, it is structured to allow daily size fluctuations, driven by changes in hepatocyte size post feeding as well as circadian regulation, on the order of 34% in mice and 10-15% in humans (6, 7). To enable this variability in size, the normal liver capsule, as is typical of visceral fascia, is thin, well innervated and contains abundant elastic fibers (1). These characteristics create a unique niche in which distinct populations of both macrophages and fibroblasts co-exist under normal physiological conditions (8).

Changes in the liver capsule in disease are yet to be well characterized, particularly in human disease. Second harmonic generation imaging (SHG) of the surface of fibrotic rat livers showed altered collagen organization, with a denser collagen network and loss of the characteristic fiber waviness. These changes were highly dependent on severity of disease and were proposed to predict the degree of liver fibrosis (9). Other rodent studies showed that, in addition to changes in the matrix, the capsular cell population changed, with mesothelial cells and fibroblasts migrating into the liver parenchyma and contributing to the myofibroblast population (10, 11). Taken as a whole, this work suggested that the capsule is an active site of pathology in rodent liver fibrosis. However, the rodent liver capsule differs significantly from the human capsule. The mouse capsule is only around 1.7 μm^2^/μm, with mesothelial cells, fibroblasts and macrophages interspersed within a very thin layer of matrix, while in contrast the normal human capsule is 30 μm–100 μm thick with mesothelial cells found on the surface and fibroblasts and macrophages embedded within a rich network of matrix (8, 10).

During the progression of liver fibrosis, collagen crosslinking as well as excessive matrix deposition leads to increasing resistance to mechanical loads, which serves as an indicator of disease severity (12, 13). Several non-invasive methods including elastography are used clinically to determine the stage of disease. In normal livers, the liver capsule is not visible under CT or MR imaging; however, this changes with disease (14), suggesting the presence of macroscopic changes, potentially including mechanical changes that could provide metrics to determine disease progression. In this study, our aim was to characterize the matrix and cellular components and accompanying mechanical properties of the capsule in human liver disease and to establish whether these are dependent on severity or etiology of disease.

## Methods

### Human samples

10 control, 6 steatosis, 7 moderate fibrosis & 37 cirrhotic anonymized human liver samples were obtained from E.E.F. at the University of Pennsylvania under institutional review board (IRB) protocol #831726 and N.D.T. at New York University under IRB protocol #il8-01106. Samples were from autopsies (controls, steatotic and cirrhotic), intraoperative biopsies taken during gallbladder or bariatric surgery (steatosis and moderate fibrosis), metastatic colon cancer resections (uninvolved tissue as controls) and liver explants (cirrhosis). Cirrhotic samples reflected a range of disease etiologies including non-alcoholic fatty liver disease (NAFLD), ethanol induced (ETOH) liver disease, primary sclerosing cholangitis (PSC), chronic hepatitis C infection (HCV), heart disease, and other causes which include autoimmune hepatitis, polycystic liver disease, hilar cholangitis and iron overload secondary to thalassemia. Due to limited numbers of sections for certain samples, a subset of randomly selected samples per group were selected for each staining. Age and sex of patients are shown in Supplemental Fig. 1. All samples were reviewed blindly by N.D.T and E.E.F. before being classed in one of four groups. Samples with no steatosis, cell infiltration or fibrosis were defined as controls, samples with a fat score of 1-3 and no fibrosis were defined as steatosis, samples with or without fat but a fibrosis score of 1-3 were defined as moderate fibrosis and samples with a fibrosis score of 4 were defined as cirrhotic samples.

### Histology and staining

Livers were cut into 1-2 cm long pieces containing the capsule layer and the underlying parenchyma, and were formalin fixed and paraffin embedded before being sectioned at a thickness of 5 μm. Before staining, sections were deparaffinized in xylene, rehydrated in a gradient of ethanol solutions and washed in diH_2_O. Next, heat-mediated antigen retrieval was carried out in 10 mM citric acid, pH6.

For diaminobenzidine (DAB) labelling, sections were incubated in 3% H_2_O_2_ to quench endogenous peroxidases (Sigma Aldrich, Rockville, MA), and then blocked for avidin/biotin (Vector Laboratories, Newark, CA) followed by StartingBlock™ T20/phosphate buffered saline (PBS) Blocking Buffer (Thermo Fisher Scientific, Waltham, MA). Samples were incubated in primary antibodies (staining buffer-0.2% Triton X-100, 0.1% bovine serum albumin, in PBS) overnight at 4°C. After washing in PBS, samples were washed and incubated with appropriate biotinylated secondary antibodies (Vector laboratories, Newark, CA) for 1 h at room temperature. Hyaluronic acid/hyaluronan (HA) was stained overnight using biotinylated HA binding protein (Millipore, Burlington, MA; 385911). All sections were washed and incubated with VECTASTAIN^®^ ABC-HRP kit (Vector laboratories, Newark, CA). Staining was developed with a DAB substrate kit (Vector laboratories, Newark, CA). Elastic fibers were stained using an elastic stain kit (Sigma-Aldrich, HT25A-1KT) per the manufacturer’s instructions. Sections were imaged by brightfield microscopy using a Nikon E600 and NIS-elements software (Nikon, Melville, NY).

For fluorescence staining, sections were blocked with StartingBlock™ T20/phosphate buffered saline (PBS) Blocking Buffer before incubation in primary antibodies (staining buffer) overnight at 4°C, followed by washing and incubation with fluorescent secondary antibody for 1 h at RT in the dark. Staining was imaged on a Zeiss Axio Observer 7 inverted microscope (Zeiss Axiocam 701 monochrome CMOS camera) and Zen blue software (Zeiss, White plains, NY). All antibodies used are listed in Supplemental Table 1.

For picrosirius red (PSR) staining, paraffin-embedded sections were dewaxed and rehydrated as above. Sections were then incubated in PSR stain (Poly Scientific R&D, Bay Shore, NY) for 1 h at room temperature. Sections were then quickly washed in acidified water (0.5% acetic acid in H_2_O), and dehydrated in 3 washes of 100% ethanol before clearing in Xylene and mounting.

### Second harmonic generation (SHG) imaging

SHG imaging was carried out on fixed sections (2D) as well as on a subset of non-processed samples (3D). Fixed and paraffin-embedded sections were imaged without dewaxing/rehydration (2D). Capsules were also carefully peeled off from the surface of fresh livers before being formalin fixed and imaged (3D). All samples were imaged using a Leica SP8 confocal/multiphoton microscope and Coherent Chameleon Vision II Ti:Sapphire laser (Leica, Buffalo Grove, IL, USA) tuned to a wavelength of 910 nm and the Leica application suite (LAX, Deerfield, Il) software. SHG of collagen generates both forward and backward signal; directionality of scatter is dependent on the size and orientation of the fibrils, with backward scatter associated with more immature fibrils (15).

### Image analysis

Thickness, percentage area and cell counting were analyzed using ImageJ (16). 3-5 images per liver were taken using the same parameters and analyzed using manually-determined ROIs. Thickness was measured manually, with 5 measurements per image. Quantification of percentage area stained was carried out using the deconvolution (H DAB) tool followed by thresholding. Cells were counted using the particle analysis tool. CD31 structures were counted manually using the count tool. Structures were defined as 2 or more clustered cells; some of these formed vessel-like structures.

PSR was analyzed under polarized light with 6-10 images analyzed per sample. The PSR dye enhances the natural birefringence of the collagen when exposed to polarized light (17). PSR images were used to estimate a uniformity index of collagen fiber distribution, as previously described (18). This index calculates values between 1 (when the objects are distributed in a regular array) and 0 (when maximal clustering occurs). In order to apply this method, a subsampled binary image of the collagen pattern was obtained by extracting pixels corresponding to the points of a superimposed regular/systematic grid.

Crimping of collagen fibers was assessed on z stacks of SHG images. Depth and length were manually measured using the line tool in ImageJ. Approximately 20 fibers were measured in 3-8 images per sample. Orientation and alignment of collagen fibers were also assessed on SHG Z-stacks using a previously described method (19).

### Mechanical testing

Human liver capsules were carefully peeled away from parenchyma using forceps before being frozen until use. Unfixed frozen samples were defrosted and placed into 1% phosphate-buffered solution for the duration of mechanical testing. Membrane cross-sectional area was calculated using a custom laser-based measurement device. This device uses a confocal displacement laser (CL-3000, Keyence Corporation), which has the capacity to measure transparent samples. Tensile testing was performed on a test frame (5542, Instron Inc.) equipped with a 10 N load cell. Samples were prepared on both ends using 400-grit sandpaper held in place with cyanoacrylate glue. Using custom aluminum grips, specimens underwent a tensile ramp-to-failure at a rate of 0.03mm/s. Outcome metrics included max load (N/mm), stiffness (N/mm), max stress (MPa), max strain (mm/mm), and modulus (MPa)(20).

### Statistical analysis

Statistical significance was assessed using one-way ANOVA followed by Tukey’s post hoc analysis, one or two-tailed student t-test or a Pearson’s correlation, calculated with Prism 9 (GraphPad Software, La Jolla, CA, USA). One-tailed student t-tests were used for difference in crimping and alignment as these were hypothesized to be reduced in cirrhotic samples. P<0.05 was regarded as statistically significant. The tests used for each analysis are noted in the figure legends.

## Results

### Capsule thickness and number of cells increase in liver disease patients

Liver capsules from control, steatotic, moderately fibrotic and cirrhotic patients were assessed for capsule thickness and cellularity. Capsules of cirrhotic samples were noticeably thicker when assessed by H&E staining, which also demonstrated that thickening was heterogeneous within a given patient sample (Fig. 1A, B). Thickening of the capsule was dependent on severity of disease but independent of etiology (Fig. 1 A-C). Quantification of thickness shows that steatotic and moderately fibrotic samples were unchanged when compared to controls; however, cirrhotic samples were approximately 3.3x thicker than controls (231.58 μm ± 21.82 and 70.12 μm ± 14.16, respectively) (Fig. 1B). Additionally, capsule thickness positively correlated with parenchymal fibrosis (Fig. 1C). Capsules from cirrhosis of different etiologies had similar ranges of thickness and no clustering was observed. Next, the number of cells found within the capsule was counted. Control and steatotic samples had similar numbers of cells (2.19×10^−4^±2.74×10^−6^ and 2.58×10^−4^±4.76×10^−6^nuclei/pixel respectively); however, both moderately fibrotic and cirrhotic had an increased number of cells (5.76×10^−4^ ± 7.26×10^−6^ and 4.157×10^−4^ ± 3.47×10^−6^ nuclei/pixel respectively) per capsule area (Fig. 1D), suggesting that an increase in cell number precedes thickening of the capsule.

**Fig. 1.**
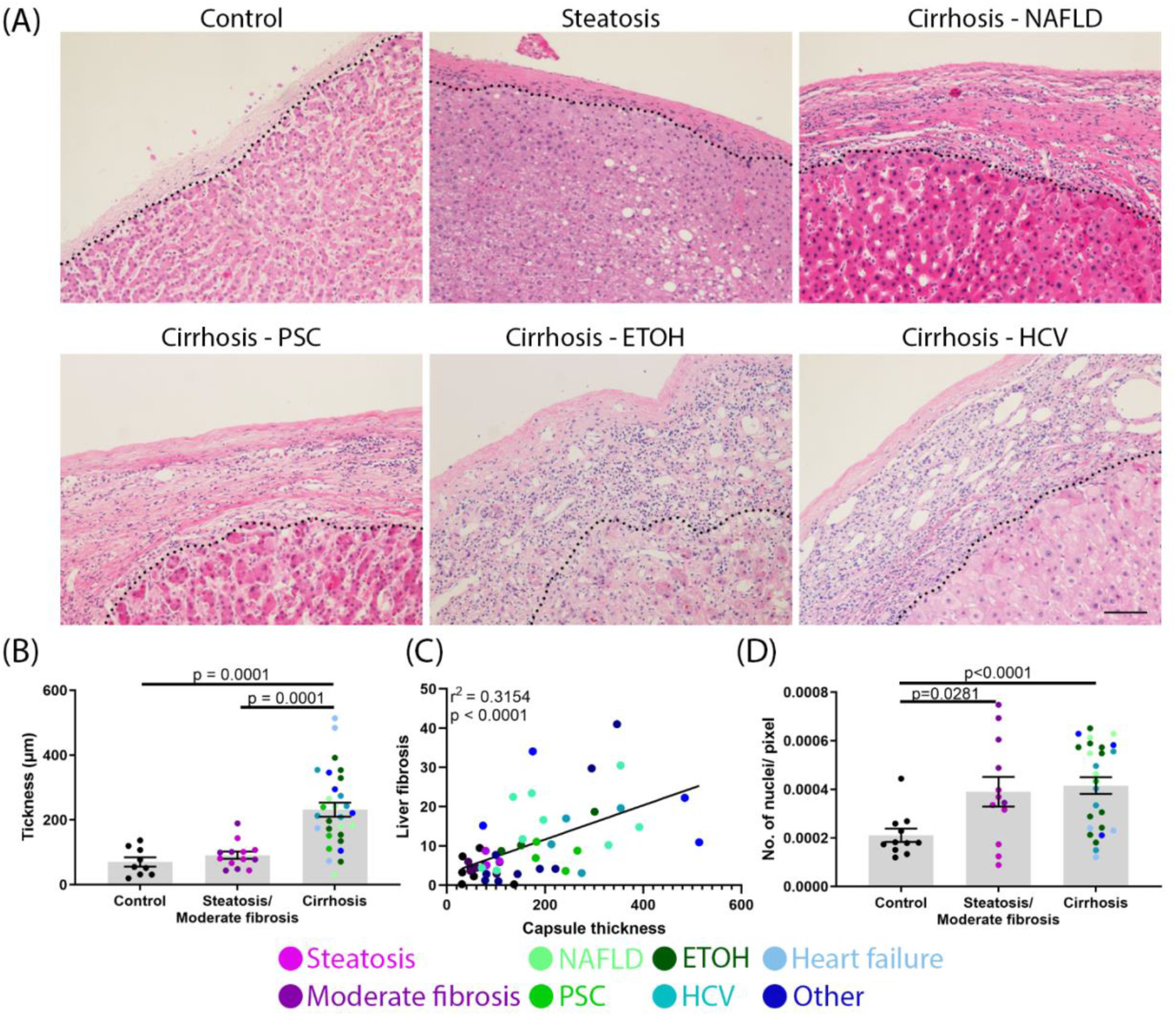
Changes in liver capsules in patients with liver disease. (A) Representative H&E images of human livers with no disease (control), steatosis and cirrhosis of different etiologies. Capsules are found above the dotted line. (B) Quantification of liver capsule thickness. (C) Correlation between parenchymal fibrosis (% collagen) and capsule thickness. (D) Quantification of cell number in the capsule. Control vs steatosis (not significant), control vs. moderate fibrosis (p=0.0004), steatosis vs. moderate fibrosis (2.578×10^−4^ ± 4.76010^−5^ and 5.762×10^−4^ ± 7.262^-5^ respectively, p=0.0052), moderate fibrosis vs. cirrhosis (NS). Thickness and number of nuclei were analyzed by one-way ANOVA and are shown as mean ± SEM. Correlation analyzed by Pearson’s correlation. N=9-31. Scale bar 50 μm.

### Increase in vascular, immune cell, fibroblast and biliary epithelial cell populations in capsules from cirrhotic livers

In order to identify the cells present in the capsule, samples were stained for vascular endothelial, immune, fibroblast and biliary epithelial cell markers. Cirrhotic capsules were richer in vascular structures (CD31+) (Fig. 2A, B) than control samples. There was also a large population of small circular leukocytes (CD45+) in the lower layers of the capsule of cirrhotic samples (Fig. 2A, C). Macrophages (CD68+) have previously been shown to be present in human capsule (8), however these did not appear to be important contributors to the immune cell population in cirrhosis (Fig. 2A).

**Fig. 2.**
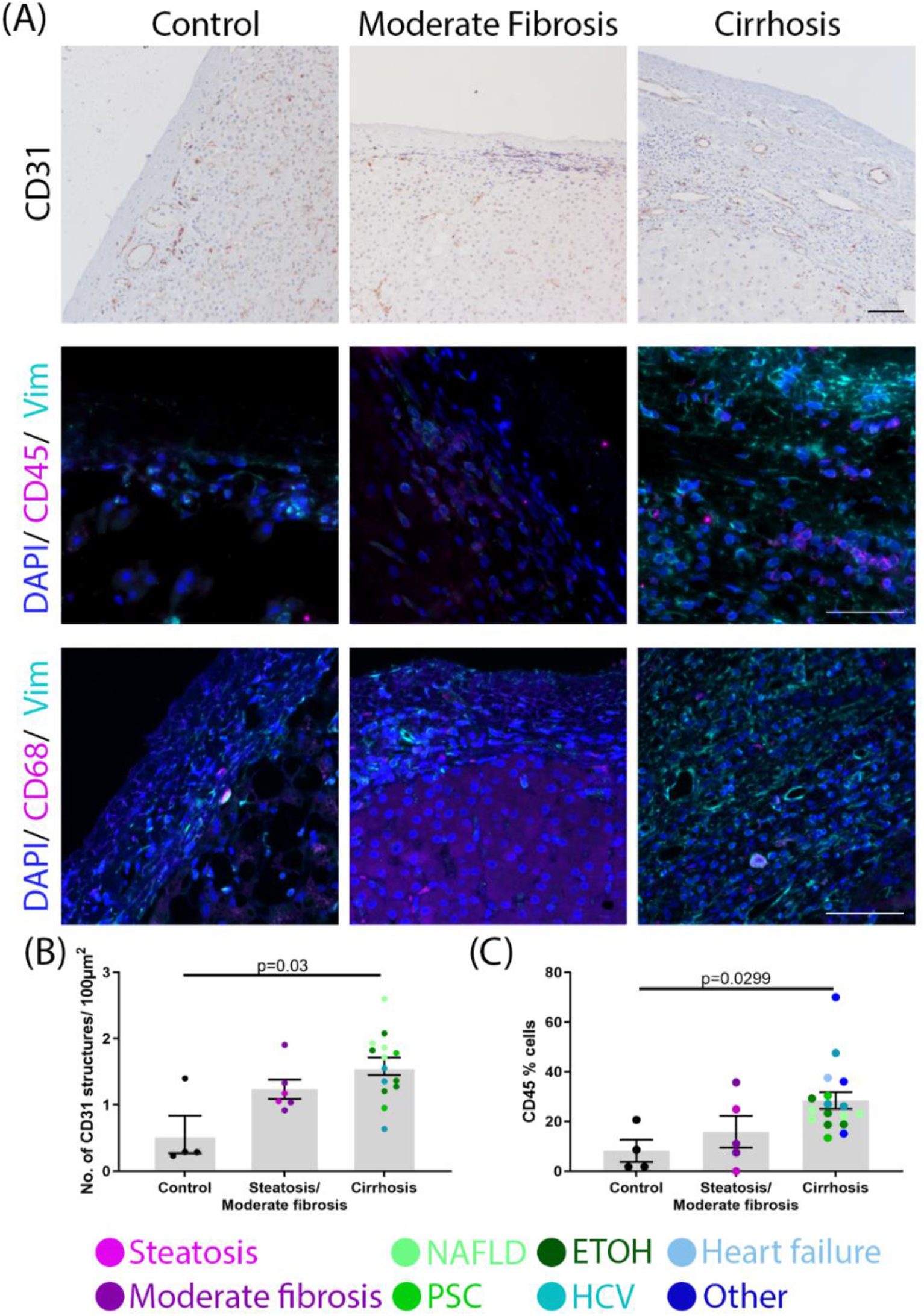
Increase in vascular and immune cells. (A) Representative images of control, moderately fibrotic and cirrhotic samples (NAFLD in this example) stained for CD31 (vascular endothelial cells; brown), DAPI (nuclei; blue), CD45 (leukocytes; pink), CD68 (macrophages; pink), or vimentin (fibroblasts; teal). (B) Quantification of the number of CD31-positive structures. (C) Quantification of CD45-positive cells in the capsule. Data were analyzed by one-way ANOVA and are shown as mean ± SEM. N=4-17. Scale bars 50 μm.

Fibroblasts, identified by vimentin staining, were found in all groups (Fig. 2A); however, activated fibroblasts (αSMA+) were significantly increased only in cirrhotic samples (Fig. 3A, B). Biliary epithelial cells (BECs) and mesothelial cells were stained using keratin 7 and 19 (Fig. 3 A, C). Mesothelial cells were found (but not consistently) on the surface of the liver capsule of all sample groups. This was confirmed by calretinin staining (Supplemental Fig. 2), in which few, if any, calretinin positive cells were observed within the capsule. Several, however, can be seen detaching from the surface, suggesting that the lack of mesothelial cells on the surface of our samples may be an artifact of tissue handling. Biliary epithelial cells (positive for keratin 7 and 19 and negative for calretinin) were found in lower layers of the capsule, often at the interface with the parenchyma. These were organized in ductal structures as well as present as individual cells and were most significantly increased in the cirrhotic patients (Fig. 3C). To determine whether cell proliferation contributed to the observed increase in cells, capsules were stained with Ki67, a nuclear protein marker for cellular proliferation. Overall there was no change in proliferation observed in the disease groups compared with controls (Fig. 3D, Supplemental Fig. 3).

**Fig. 3.**
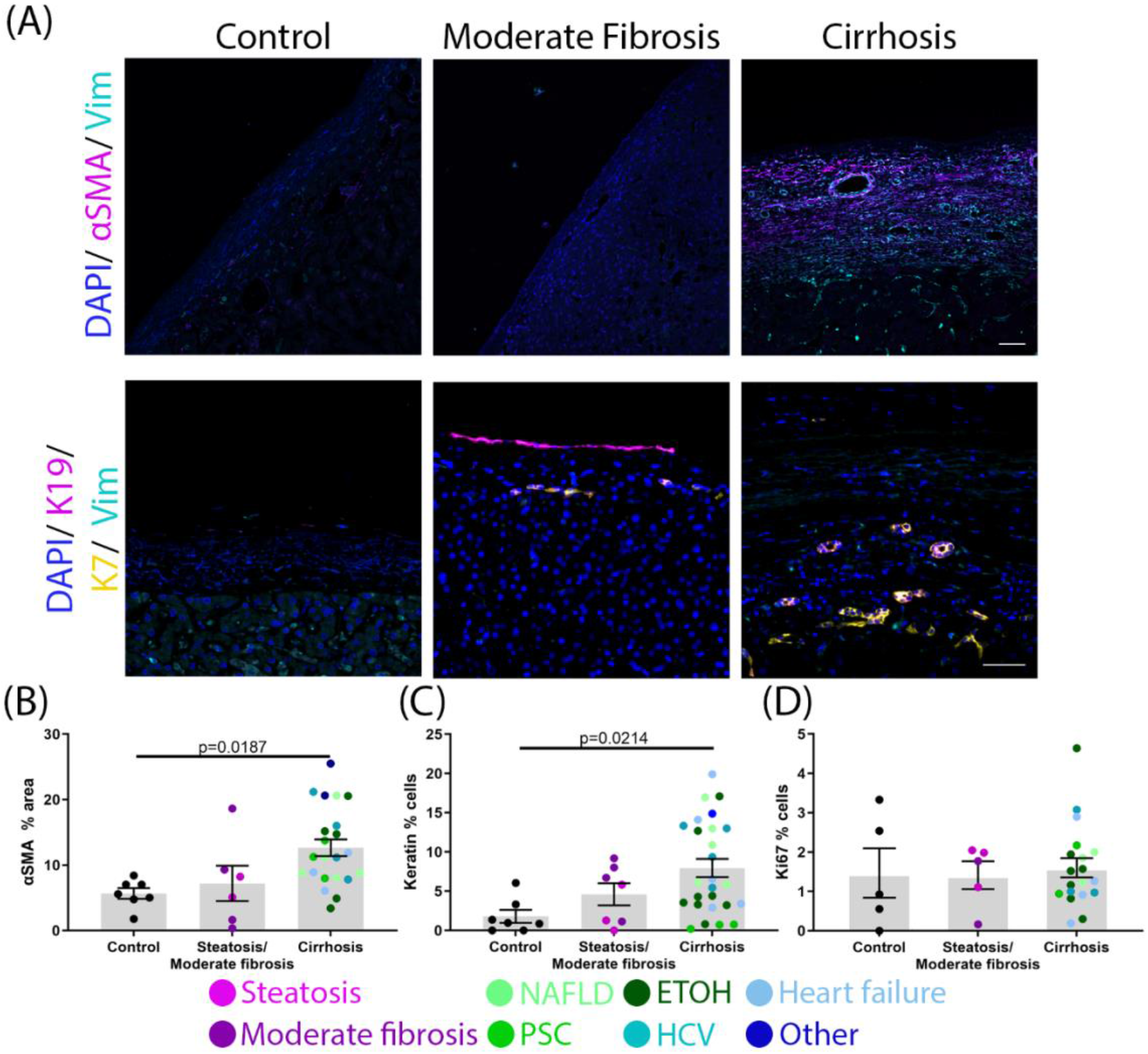
Increase in fibroblasts and biliary epithelial cells in diseased capsule. (A) Representative images of control, moderately fibrotic and cirrhotic samples (NAFLD in this example) stained for DAPI (nucleus; blue), vimentin (fibroblasts; cyan), αSMA (activated fibroblasts; magenta), keratin 19 (K19; magenta), and keratin 7 (K7; yellow). (B) Quantification of αSMA percentage area. (C) Quantification of keratin-positive cells (pink) in the capsule. (D). (E) Quantification of Ki67-positive cells in the capsule. Data were analyzed by one-way ANOVA and are shown as mean ± SEM. N=5-24. Scale bars 50 μm.

### Collagen fibers and organization are altered in patients with liver cirrhosis

Collagen fibers and organization were visualized by SHG microscopy (Fig. 4). There appeared to be two layers (the upper 70 μm and the remaining lower layer), and the content of collagen was quantified in each. The percentage area of collagen was consistent for the top layer for all disease states; however, capsules from cirrhotic samples had significantly less dense collagen in the lower layer (Fig. 4A, B). We calculated the uniformity index (which quantifies regularity of collagen fibers) from 2D SHG images and found that collagen fibers in control and steatotic/moderately fibrotic samples were distributed in a relatively uniform manner, while collagen fibers in cirrhotic capsules were more irregular (Fig. 4A, C).

**Fig. 4.**
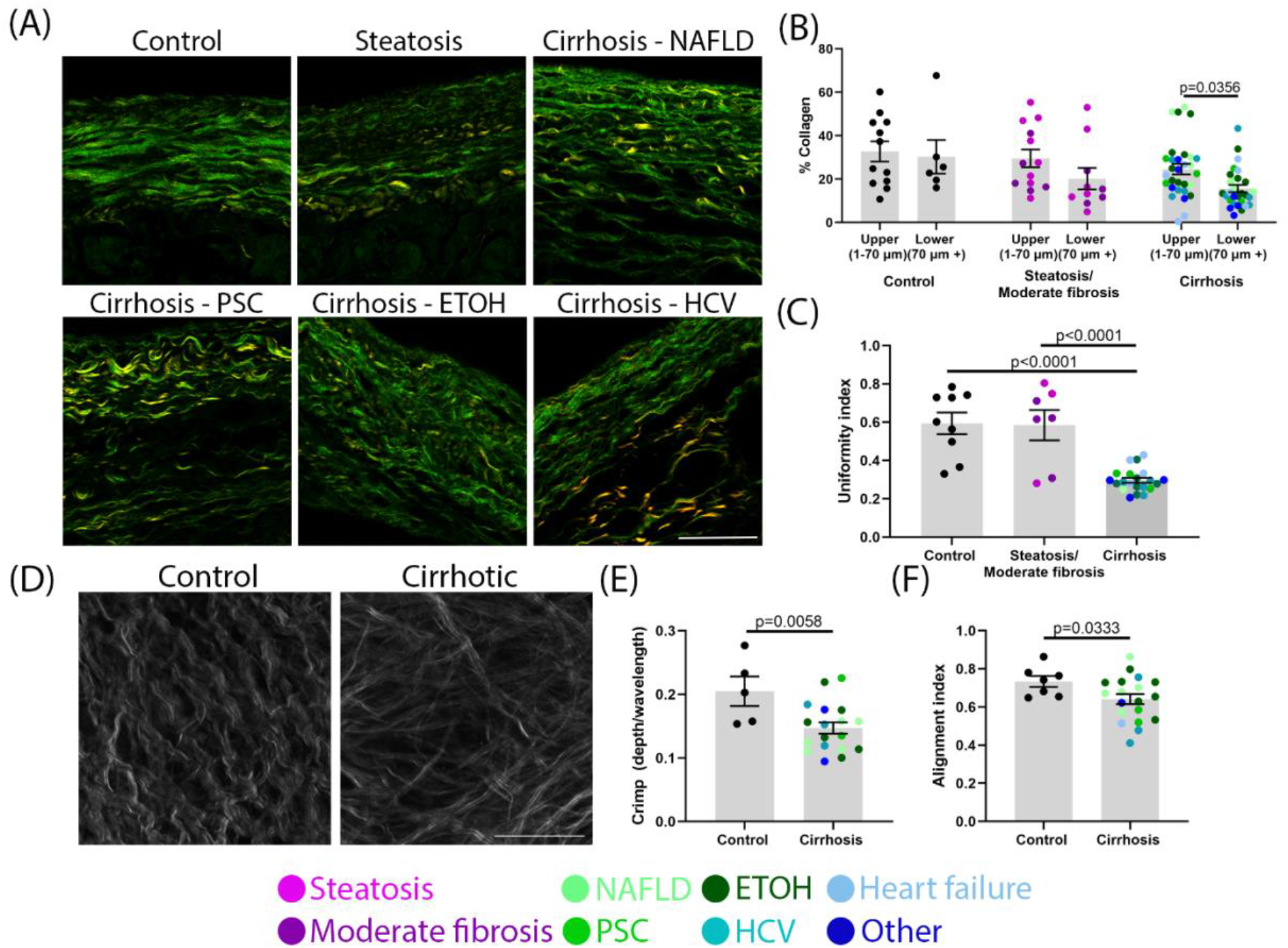
Organization of the capsular collagen network is disrupted in cirrhosis. (A) Representative SHG images of cross sections of control, steatotic, or cirrhotic samples. (B) Quantification of percentage area of collagen in the upper layer and lower layer of capsule. N=12-31. (C) Calculated uniformity index. N=7-20. Data were analyzed by one-way ANOVA and are shown as mean ± SEM. (D) Representative top-down SHG images of a control and a cirrhotic sample. (E) Calculated collagen fiber crimp. (F) Calculated alignment index of collagen fibers. Data analyzed with an unpaired one-tailed student t test and shown as mean ± SEM. N=5-18. Scale bar 50 μm.

To further understand the organization of collagen in the capsule, 3D top-down SHG imaging was done on freshly fixed samples (Fig. 4D). Individual collagen fibers in control samples had a characteristic crimp, which is often found in healthy tissue (21) and which provides low levels of absorption and compliance. Collagen fibers in cirrhotic samples largely lost this crimp although bundling of fibers, as assessed by SHG imaging, appeared unaffected (Fig. 4D, E, Supplemental Fig. 4). Collagen fibers in controls were organized in highly aligned sheets layered perpendicularly on top of each other creating a 3D cross hatch pattern that has also been described as a “two fiber family” organization (22). This was quantified using a previously described method (19) which assigns fibers to directional families, clearly showing that in most cases two individual directional families alternate in contributing to each layer of the collagen network (Supplemental Fig. 5). The alignment index, which measures how aligned fibers are within these assigned families, was considerably reduced, suggesting that overall organization is disrupted in capsules from cirrhotic livers (Fig. 4D, F).

### ECM composition of liver capsule is altered in patients with liver cirrhosis

Liver fibrosis leads to the excessive deposition of a complex network of structural and biochemically-active matrix proteins in the liver parenchyma that in turn help drive progression of disease. The ECM composition of the capsule, however, and how this changes in disease is unclear. We used immunostaining to define changes in matrix composition. We found layers of thick elastic fibers, important for capsule stretch and recoil during fluctuations in liver size, in the control samples. Scattered HA was identified in the lower layer of the healthy capsule. Collagen 3, important for collagen fibrillogenesis as well as repair and regeneration of tissues (23) is also found in relatively abundant amounts throughout the capsule. Versican, a proteoglycan, was found in limited amounts. Capsules from steatotic and moderately fibrotic livers showed similar distributions of these matrix components (Fig. 5A-E). In cirrhosis, elastic fibers and collagen 3 were increased but in proportion to the increase in thickness, leading to a similar overall concentration (% area) and collagen 1:3 ratio as in controls (Fig. 5B, D, Supplemental Fig. 6). Both HA (control 13.52 ± 5.261 % vs cirrhosis 35.99 ± 3.505 %) and versican (control 0.6797 ± 0.4168 % vs cirrhosis 17.50 ± 3.010 %) were significantly increased, primarily in the lower capsule at the interface with the parenchyma (Fig. 5A, C, E).

**Fig. 5.**
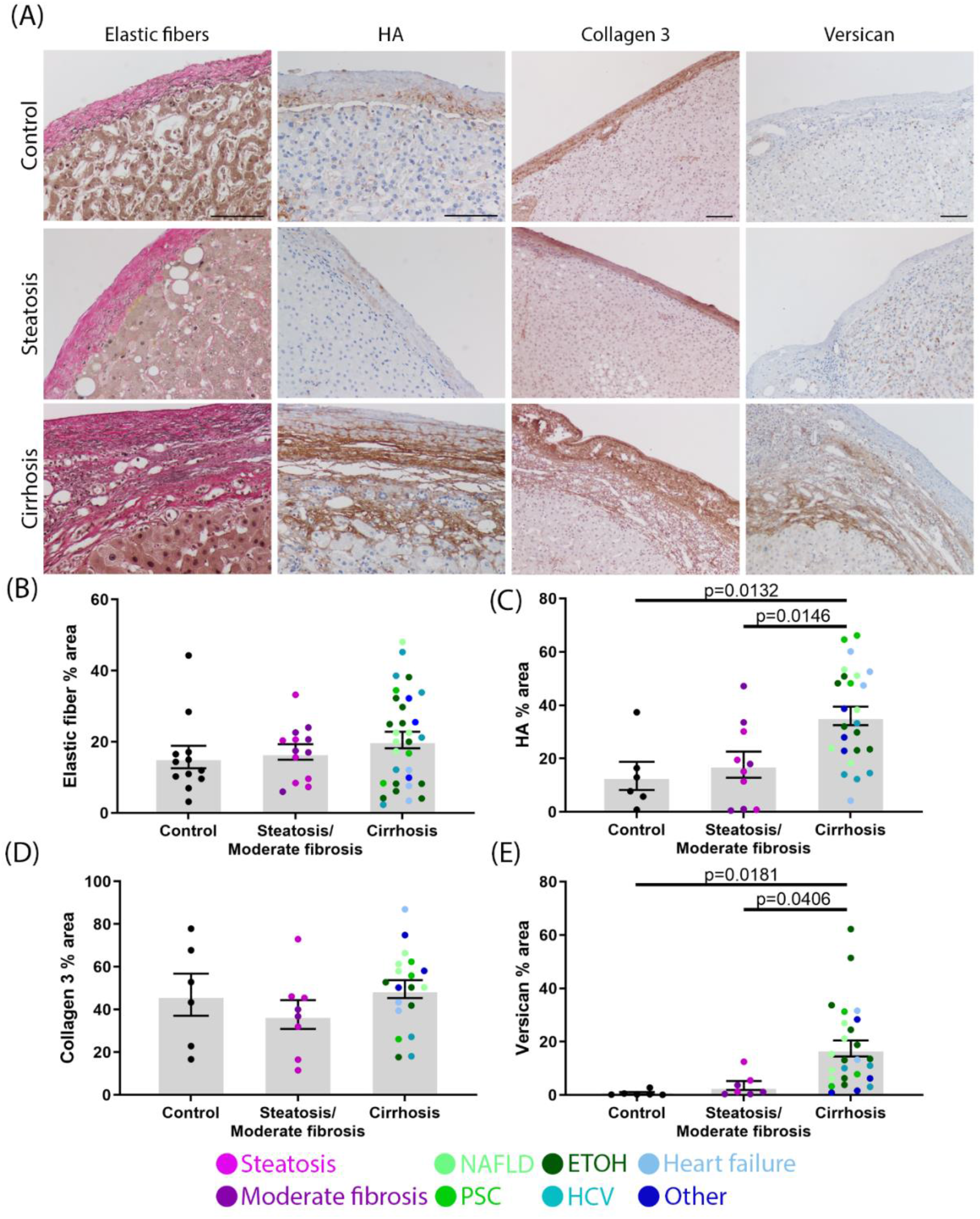
Deposition of key matrix components in the liver capsule. (A) Representative imaging of elastic fiber (Van Geison stain; black), HA (brown), collagen 3 (brown) and versican (brown) staining from patient livers with no visible disease (control), steatosis or cirrhosis (NAFLD in this example). (B) Quantification of elastic fiber percentage area. N=12-31. (C) Quantification of HA percentage area. N=6-26. (D) Quantification of collagen 3 percentage area. N=6-19. (E) Quantification of versican percentage area. N=7-26. Data were analyzed by one-way ANOVA and are shown as mean ± SEM. Scale bar 100 μm.

### Cirrhotic liver capsule is less resistant to mechanical load

To establish whether the observed changes in the composition and organization of the capsular matrix affect its function, we tested capsule mechanical properties. Cross-sectional areas of peeled capsules were measured and specimens were placed on a test frame (Fig. 6A) where force-displacement profiles were recorded (Fig. 6B). Several mechanical properties of interest were calculated, including stiffness (N/mm), which is the slope of the force-displacement curve. Additionally, stress, which is the amount of force exerted on the tissue divided by cross-sectional area, and strain, which is the amount of displacement of the tissue relative to its initial length, were calculated. Normalized measures of strength were calculated from the stress-strain curve; including the elastic modulus (MPa), which is the slope of the stress-strain curve. Clear clustering of control and cirrhotic capsules can be seen (Fig. 6C), suggesting altered mechanics in disease. The liver capsule is known to have biphasic mechanics, with an initial linear phase at low strain (0-7%) and a second linear phase at higher strain (10 to 30%) (24, 25). Unexpectedly, cirrhotic capsules required less force and stress to fail than controls (Fig. 6D, E), and had decreased moduli (Fig. 6F, Supplementary Fig. 7B). The moduli of cirrhotic capsules were significantly less than control capsules in both the toe region (low strains; 13.42 ± 2.61 and 34.11 ± 11.92 MPa, respectively) and the elastic region (high strains; 38.04 ± 9.47 and 123.9 ± 11.63 MPa, respectively) (Fig. 6F, Supplementary Fig. 7C).

**Fig. 6.**
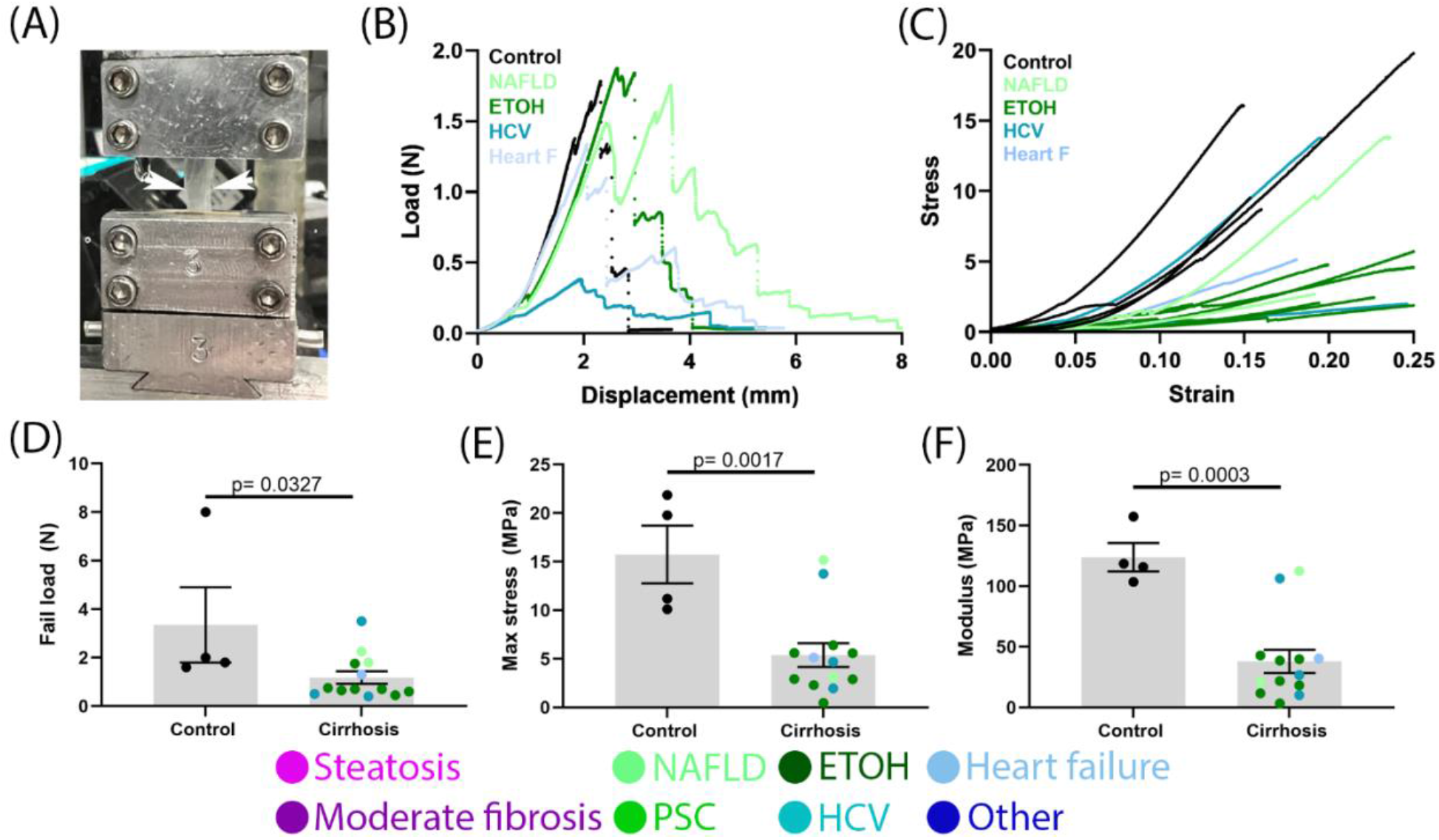
Capsules from cirrhotic patients are less stiff than those from control patients. (A) Capsule (found between two arrows) gripped in place during a tensile test. (B) Representative force-displacement curves. (C) Representative stress-strain curves. (D) Ultimate loads at tissue failure. (E) Calculated maximum stresses at tissue failure. (F) Calculated moduli at high strain. Data were analyzed with an unpaired two-way student t test and are shown as mean ± SEM. N=4-13.

## Discussion

The contribution of the capsule to the pathogenesis of liver fibrosis has not been well studied, particularly in human disease. Here we show that the thickness, extracellular matrix content and organization, cellular composition, and mechanical properties of the human capsule change significantly in cirrhosis, suggesting that the capsule is an active site of human liver disease.

The capsule is part of the visceral fascia (1); our data are consistent with published data showing that fascia plays a role in disease pathogenesis, and that changes in elastic fibers, HA and collagen, with associated fascial thickening, can lead to abnormal mechanics and function (26-31). Although strong similarities exist, exact changes in ECM composition at different sites of fascia are likely site specific.

There was a particularly significant increase in deposition of HA in the cirrhotic capsule. This is consistent with increases in HA observed in the parenchyma and, notably, in the serum of patients with cirrhosis, in whom it is used as a biomarker for diagnosis (32, 33). Increases in HA in the liver result from both increased production and decreased clearance by damaged liver sinusoidal endothelial cells (SECs) (34). In the capsule, where the number of fibroblasts is increased and there are few SECs, increased production is most likely, although decreased clearance from SECs adjacent to the capsule could also contribute. The increase in HA would almost certainly have both physical and biochemical consequences. HA undergoes swelling, creating and occupying space within a tissue, allowing for better cell migration and collagen deposition (35-37); indeed, we observed HA protruding between pairs of hepatocytes at the interface of the newly deposited capsule and parenchyma. In liver fibrosis, HA mediates hepatic stellate cell myofibroblastic activation, angiogenesis and biliary epithelial cell proliferation (38-40). Co-localization of these in the lower capsule suggests a similar effect on cells within the capsule. Versican, an HA-binding proteoglycan found primarily in the cirrhotic capsule, is also involved in collagen fibrillogenesis, cell adhesion and migration, and signaling (41, 42). It is a likely contributor to the change in the collagen network organization observed within the capsule, given that it is known to bind collagen fibers directly and to drive collagen bundle organization (41, 43). Versican may also induce a pro-fibrotic phenotype in capsular fibroblasts and adjacent hepatic stellate cells (42, 44).

Our data show that the majority of changes in the capsule in fibrosis occur in end-stage cirrhotic patients, suggesting that capsule changes are secondary to parenchymal disease. The late manifestations of capsular disease may also reflect the unique subpopulations of both macrophages and fibroblasts residing in capsule, in particular capsular macrophages, which provide immunosurveillance against pathogens within the peritoneum and may protect against early disease (8, 45). Capsular fibroblasts also highly express the transcription factor Odd-Skipped Related Transcription Factor 1 (Osr1)(8), which is protective against inflammation and early fibrosis in a fatty liver disease mouse model (46). Thus, capsular fibroblasts as well as macrophages may have a basal anti-inflammatory phenotype.

Mesothelial cells have previously been shown in rodents to migrate into the liver parenchyma and differentiate into myofibroblasts participating in collagen deposition (11). No mesothelial cells could be observed within the capsule or in the “active zone” in our samples, suggesting that that transmigration of these cells in human disease may be limited. However, mesothelial cells were not found consistently on the surface of samples, and appeared to have detached during handling of the tissue, making any comment on this process difficult.

Mechanical testing of patient capsules showed that liver cirrhosis leads to important functional changes in the capsule. In contrast to the rest of the liver, which stiffens in fibrosis and cirrhosis, capsules from cirrhotic livers were less resistant to mechanical loads than controls (12, 13). We speculate that this is protective and a compensatory mechanism in the context of pathologically high intrahepatic pressures, preventing the further increases in pressure that would occur with a more rigid capsule. The key difference in mechanics between the liver and the capsule in disease reflects the starting mechanical properties and composition of the tissue. The bulk of the liver is highly cellular, with limited matrix, and is relatively soft. In fibrosis, the matrix is cross-linked and becomes highly organized within the tissue, leading to increased overall rigidity. In contrast, we have shown here that the normal liver capsule is a matrix-rich tissue organized into a two-fiber family pattern, consistent with measurements showing it to be a relatively stiff tissue (22). In fibrosis, key matrix properties such as crimping and organization are altered, and this likely leads to the observed softening. The crimp found in collagen fibers allows for gradual fiber extension during tensile testing, consistent with the biphasic response observed in normal liver capsule, similar to that observed in tendon (47). Collagen crimping is reduced in cirrhotic capsules, leading to decreased elasticity and increased brittleness, as observed by the lower fail load of the cirrhotic capsule compared to the control. Organization of the collagen network drives tissue stiffness – a disorganized network is less stiff than a network in which fibers are well aligned (22, 48). Previous studies have found the normal capsule to have a modulus between 27.7 MPa (24) and 48.7 MPa (25). Control samples we report here had a modulus of 123.9 MPa, which, although slightly higher, is a similar order of magnitude. There are two factors that could contribute to the higher modulus we observed. The two previously-published studies were carried out on livers from cadavers in which degradation of the tissues is likely to have started, leading to reduced tissue integrity. When comparing fresh (tested within 5 days of death) and frozen tissue, the fresh tissue appeared to have a relative decrease in modulus (16.9 MPa vs 27.7 MPa) (24). Second, the controls used in our study here are from uninvolved colon cancer metastasis, and although histology of these was normal, there may have been underlying mechanical changes.

Our data support viewing the liver capsule, a part of the interstitium and visceral fascia, as an active albeit secondary site of liver disease. It should be considered when developing diagnostic and predictive approaches for liver fibrosis and cirrhosis. Additionally, we note that cirrhosis leads to significant mechanical changes, resulting in a more compliant capsule that may serve as a limited compensatory mechanism to reduce portal hypertension in cirrhosis.

**Table 1:** Antibodies used.

| Primary antibodies |  |  |  |
| --- | --- | --- | --- |
| Antigen | Species | Dilution | Source |
| αSMA | Mouse | 1:100 | Sigma, A2547 |
| Collagen III | Goat | 1:100 | Southern biotech ,1330-01 |
| Cytokeratin 7 | Rabbit | 1:200 | Abcam, ab68459 |
| Cytokeratin 19 | Mouse | 1:200 | Abcam, ab7754 |
| Ki67 | Rabbit | 1:100 | Abcam, ab16667 |
| CD45 | Rat | 1:50 | Novus, NB100-77417SS |
| Vimentin | Chicken | 1:500 | Novus, NB300 223 |
| CD68 | Mouse | 1:100 | Abcam, ab955 |
| CD31 | Rabbit | 1:100 | Novus, NB100-2284 |
| Calretinin | Rabbit | 1:100 | Thermofisher, PA5-32287 |
| Versican | Rabbit | 1:200 | Novus, NBP1-85432 |
| Biotinylated anti-rabbit | Donkey | 1:200 | Vector laboratories, BA-1000 |
| Biotinylated anti-goat | Donkey | 1:200 | Vector laboratories, BA-9500 |
| Cy 2 anti-mouse | Donkey | 1:500 | Vector laboratories ,715-225-150 |
| Cy 3 anti-rabbit | Donkey | 1:500 | Vector laboratories, 711-165-152 |
| Cy3 anti-rat | Donkey | 1:500 | Vector laboratories, 712-165-153 |
| Cy5 anti-chicken | Donkey | 1:500 | Vector laboratories, 703-605-155 |

## Abbreviations

EHBD: extrahepatic bile duct
SMA: smooth muscle actin
DAPI: 4’,6-diamidino-2-phenylindole
SHG: second harmonic generation
HA: hyaluronic acid/hyaluronan

## Acknowledgments

We would like to express our gratitude and acknowledge the Molecular Pathology and Imaging Core of the UPenn NIDDK Center for Molecular Studies in Digestive and Liver Diseases (NIH P30 DK050306), Gordon Ruthel and the Penn Vet Imaging Core (NIH S10 OD021633-01) and the Perelman School of Medicine Cell and Developmental Biology Microscopy Core for their assistance throughout this project.

---

**Supplemental Fig. 1.**
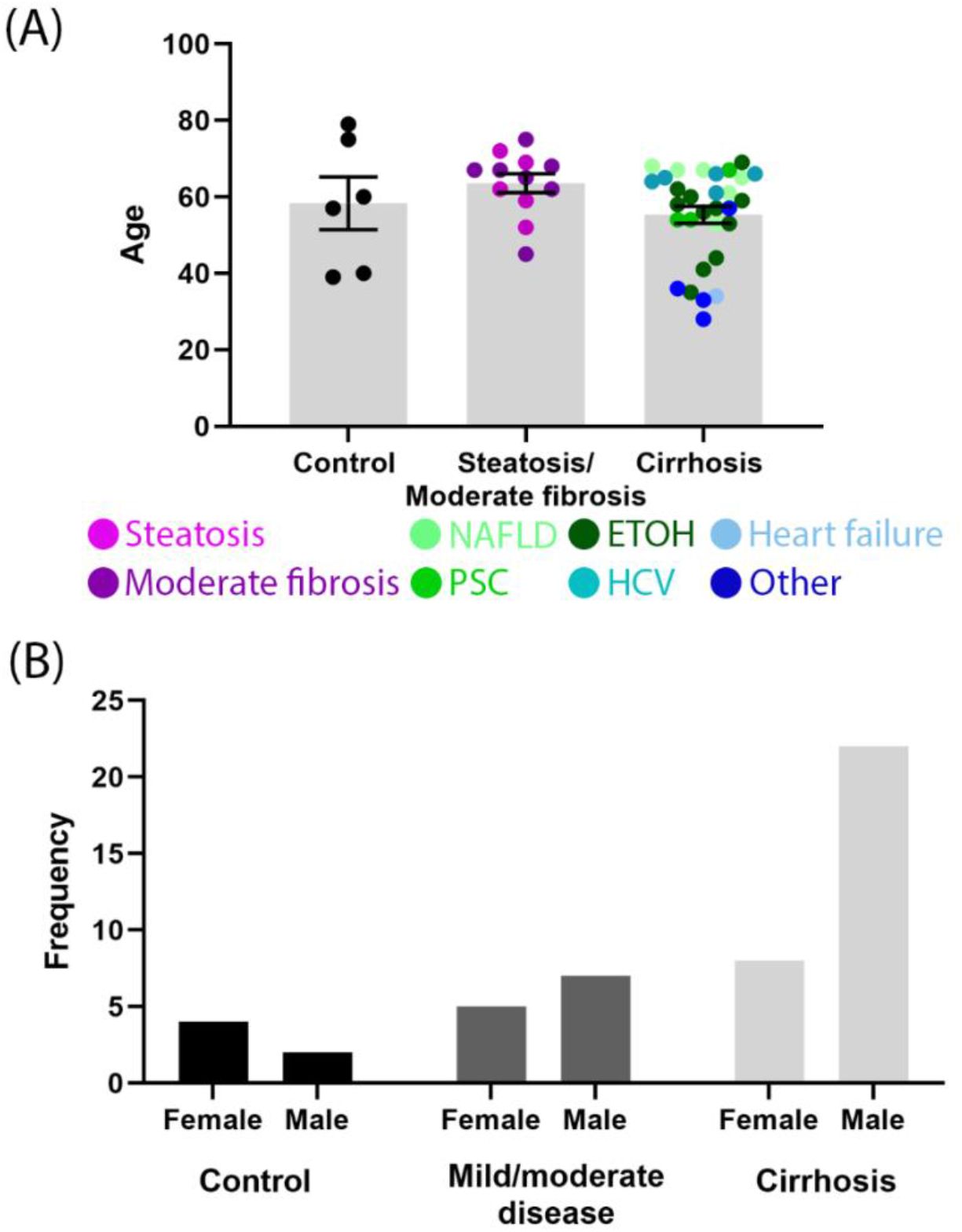
Sex and age of patient samples. (A) Patient ages for the control, streatosis/moderate fibrosis and cirrhotic samples used. (B) Sex distribution of patients for the control, streatosis/moderate fibrosis and cirrhotic samples used.

**Supplemental Fig. 2.**
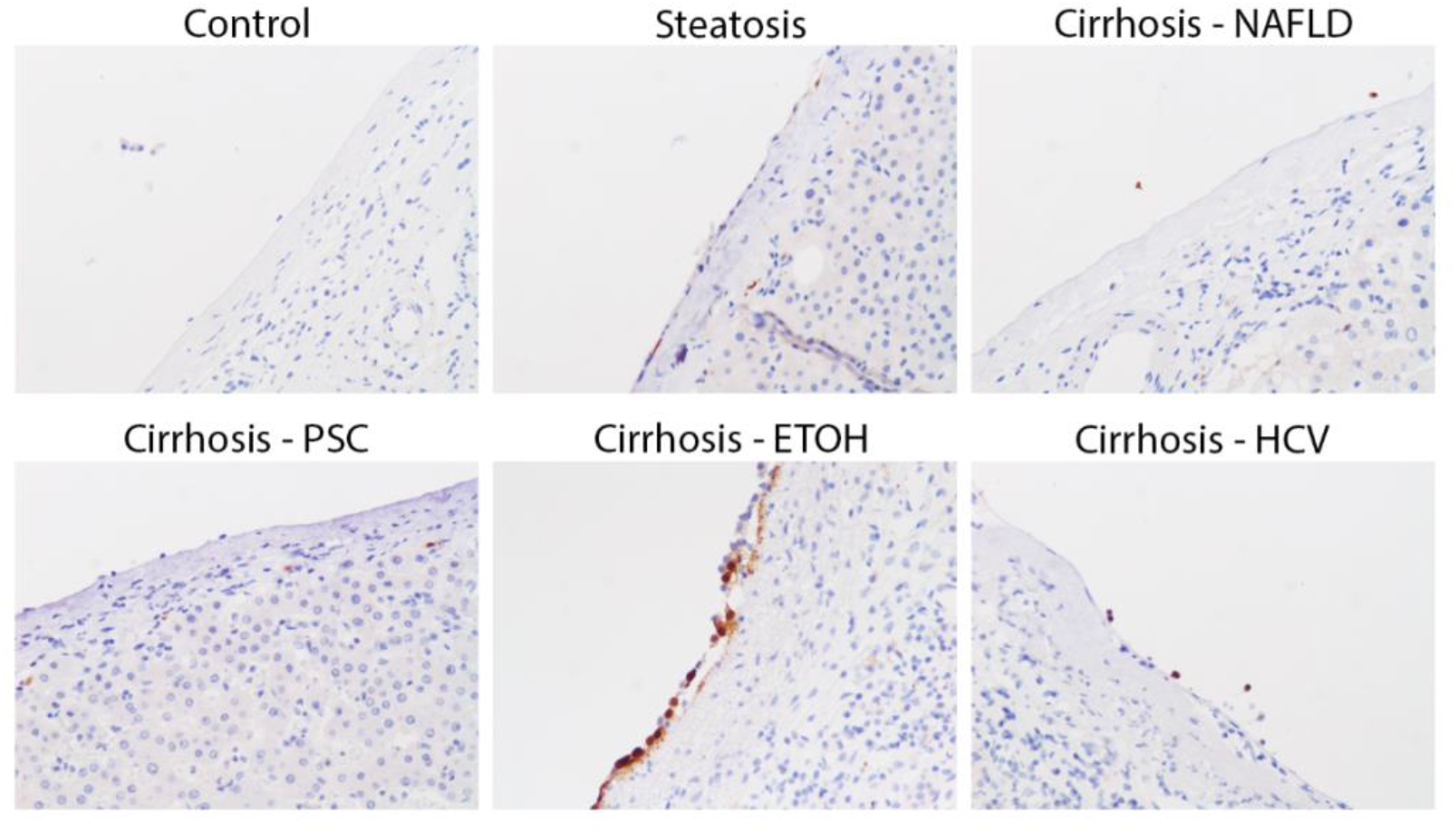
Mesothelial cells are found sporadically on the capsule surface. Representative staining for calretinin in control and steatotic samples as well as cirrhotic samples of different etiologies.

**Supplemental Fig. 3.**
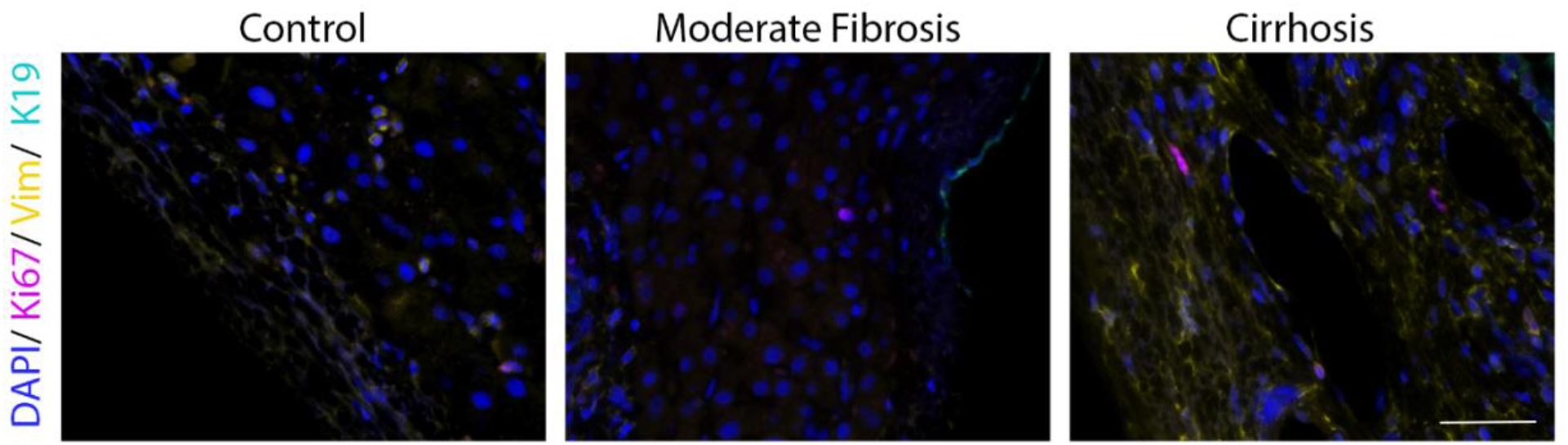
Proliferation is unchanged in disease. Representative images of control, moderately fibrotic and cirrhotic samples (NAFLD in this example) stained for DAPI (nucleus; blue), Ki67 (proliferation; magenta), vimentin (fibroblasts; yellow), and K19 (BECs; cyan). Scale bars 50 μm.

**Supplemental Fig. 4.**
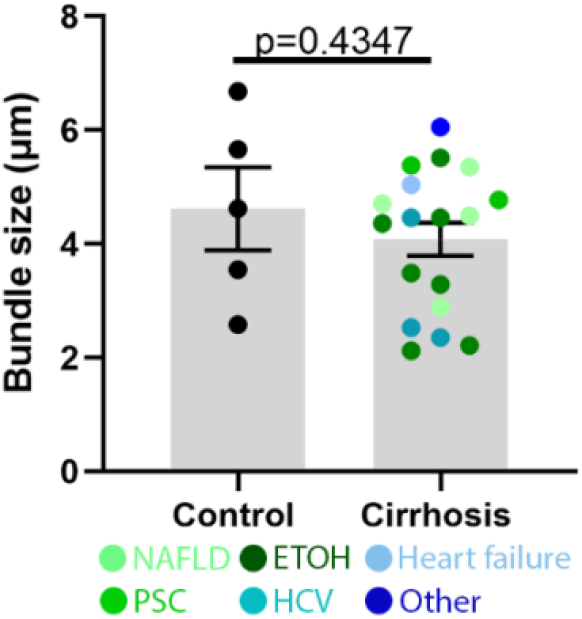
Collagen bundling is unaffected by disease. Quantification of collagen bundle size. Data were analyzed with an unpaired two-tailed student t test and are shown as mean ± SEM. N=4-17

**Supplemental Fig. 5.**
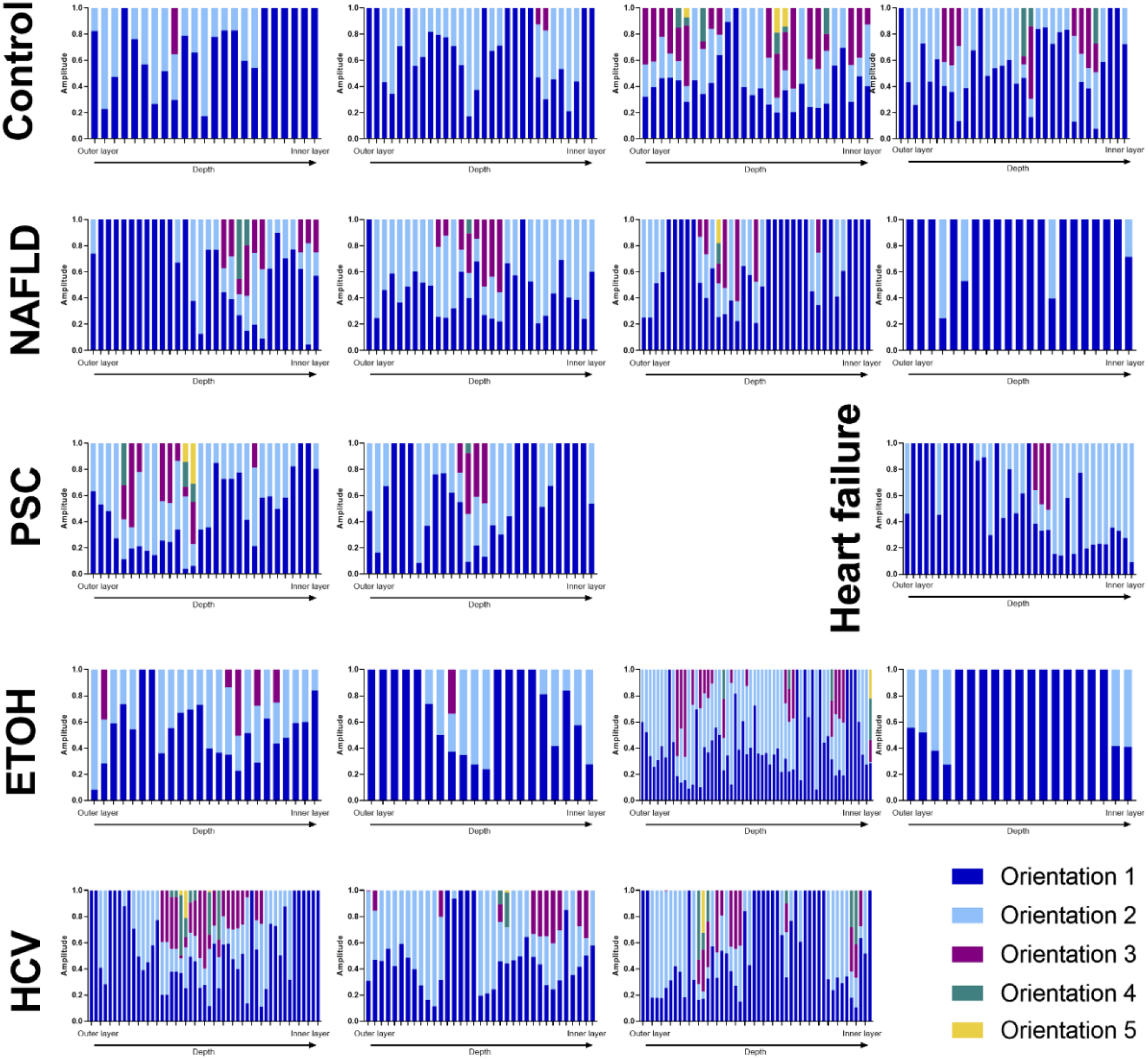
Amplitude of collagen orientation families. Fibers from SHG z stacks placed into orientation families of similar orientation for control and cirrhotic liver samples of different etiology.

**Supplemental Fig. 6.**
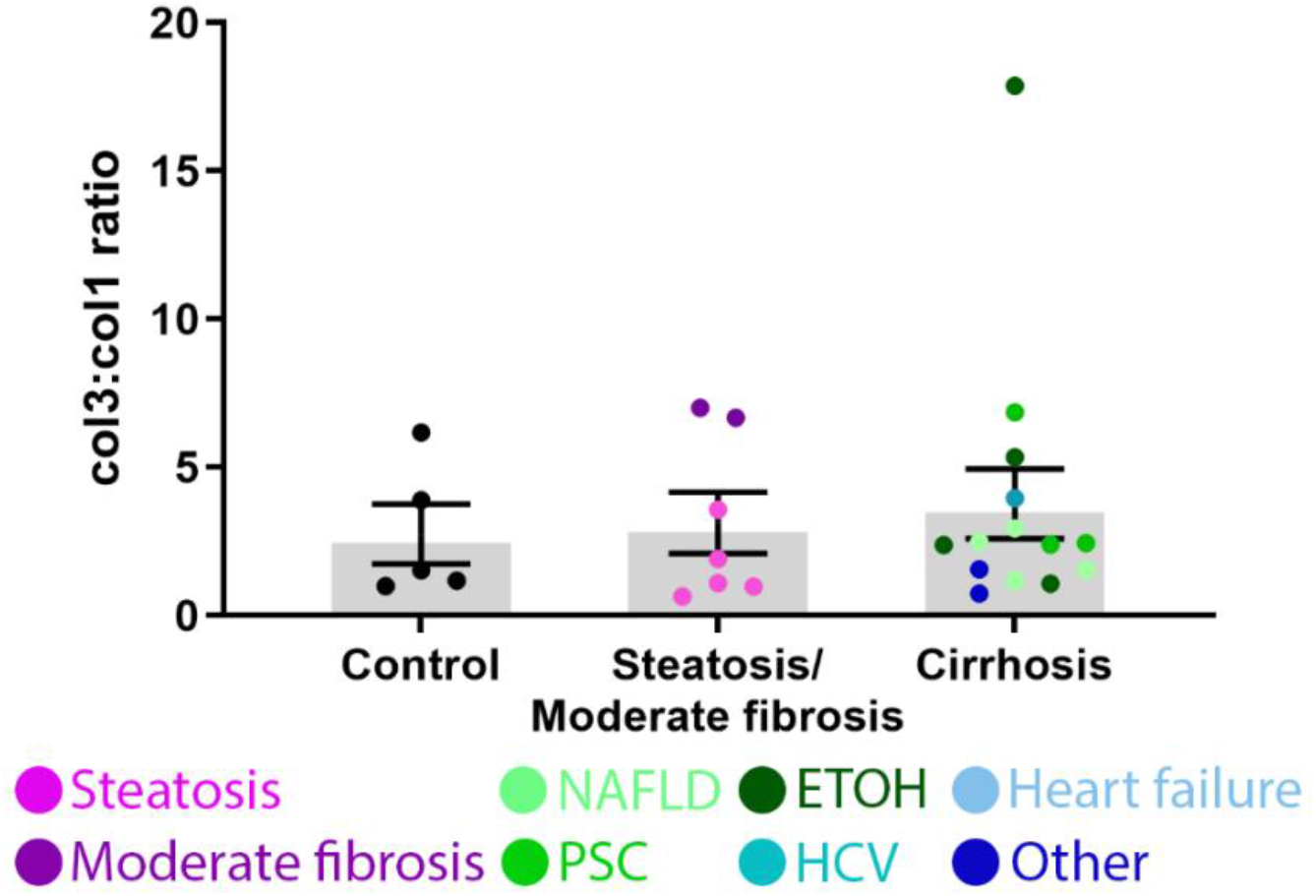
Collagen 3 to collagen 1 ratio is unaltered. Quantification of collagen 3 to 1 ratio. Data were analyzed by one-way ANOVA and are shown as mean ± SEM. N=5-14.

**Supplemental Fig. 7.**
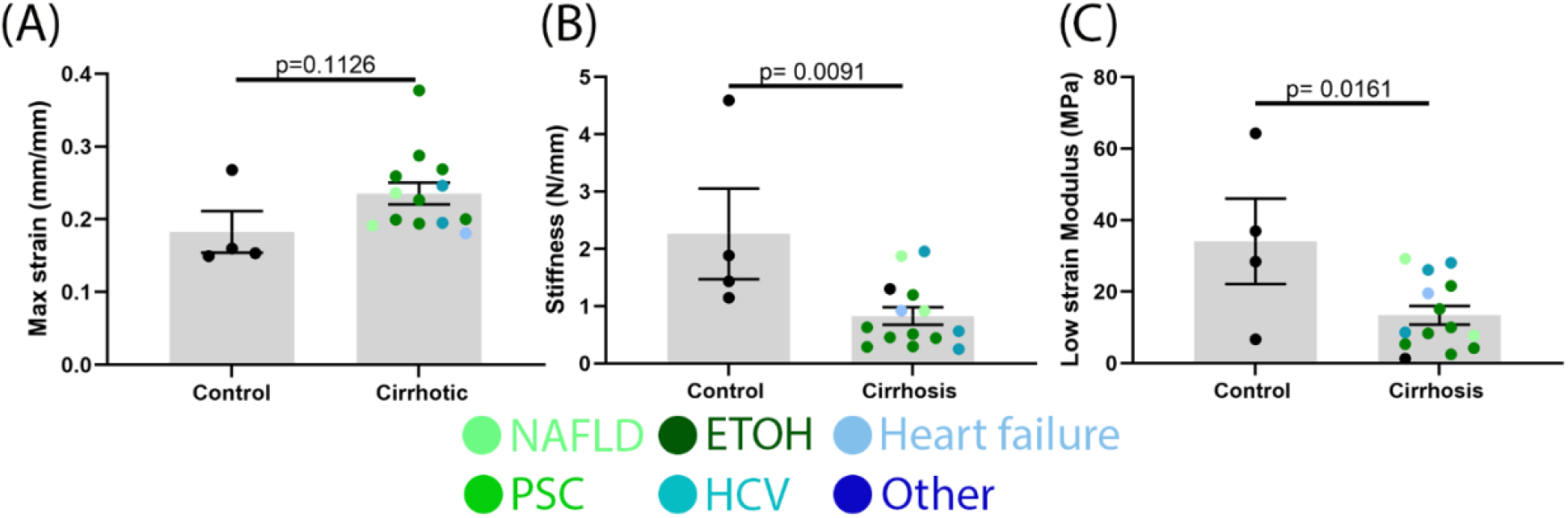
Liver capsule mechanics are altered in cirrhotic patients. (A) Maximum strain at which tissue starts to fail. (B) Calculated stiffness. (C) Calculated modulus at low strain (approx. 4-6%). Data were analyzed with an unpaired two-way student t test and are shown as mean ± SEM. N=4-14.

